# Tuning Collective Behaviour in Zebrafish with Genetic Modification

**DOI:** 10.1101/2024.04.02.587671

**Authors:** Yushi Yang, Abdelwahab Kawafi, Qiao Tong, Chrissy L. Hammond, Erika Kague, C. Patrick Royall

## Abstract

Zebrafish collective behaviour is widely used to assess their physical and mental state, serving as a valuable tool to assess the impact of ageing, disease genetics, and the effect of drugs. The essence of these macroscopic phenomena can be represented by active matter models, where the individuals are abstracted as interactive self-propelling agents. The behaviour of these agents depends on a set of parameters in a manner reminiscent of those between the constituents of physical systems. In a few cases, the system may be controlled at the level of the individual constituents such as the interactions between colloidal particles, or the enzymatic behaviour of *de novo* proteins. Usually, however, while the collective behaviour may be influenced by environmental factors, it typically cannot be changed at will. Here, we challenge this scenario in a biological context by genetically modifying zebrafish. We thus demonstrate the potential of genetic modification in the context of controlling the collective behaviour of biological active matter systems at the level of the constituents, rather than externally. In particular, we probe the effect of the lack of *col11a2* gene in zebrafish, which causes the early onset of osteoarthritis. The resulting *col11a2 -/-* zebrafish exhibited compromised vertebral column properties, bent their body less while swimming, and took longer to change their orientations. Surprisingly, a group of 25 mutant fish exhibited more orderly collective motion than the wildtype. We show that the collective behaviour of wildtype and *col11a2 -/-* zebrafish are captured with a simple active matter model, in which the mutant fish are modelled by self–propelling agents with a higher orientational noise on average. In this way, we demonstrate the possibility of tuning a biological system, changing the state space it occupies when interpreted with a simple active matter model.

## INTRODUCTION

Animal groups exhibit a rich variety of collective behaviour, exhibiting complex patterns across different length scales, from a swarm of insects (centimeters) to large fish shoals (kilometers) [1–6]. Such collective motion may be interpreted as an emergent phenomena originating from interactions between individuals [7–9], which can be beneficial for the entire group [10–14]. Therefore, a reductionist way to approach such collective behavior is to treat the individuals as interacting self-propelling agents [15, 16]. In this sense, the animal group can be described as an *active matter* system [17–19]. Particular examples include flocking and schooling behavior of biological systems which can be reproduced in computer simulations [20–22], experiments with active colloids [16, 23, 24], as well as robotic swarms [25, 26]. Within the framework of a system comprised of many agents, or particles in the case of physical systems, the macroscopic behavior of the system can be controlled externally. In physical systems, state variables such as temperature drive phase transitions, or noise in agent trajectories leads to a dynamical ordering transition in active systems [15, 27, 28]. In biological systems collective behavior in schools of fish, for example, can be strongly influenced by predation [29] and swarms of midges may be controlled ambient lighting [6, 30].

What these diverse examples have in common is a system of individuals with certain characteristics which respond to an external change in the environment. However, another *bottom-up* approach is to tune the properties of each constituent. While often this is forbidden by nature (the immutable laws of quantum mechanics in the case of atoms) or impractical (it is hard to control large bird flocks for example [9]), some systems are amenable to the design of their constituents. For example, synthetic active colloids [16] are an idealised playground to understand the principles of active matter, exhibiting dynamical [24, 31, 32] or structural [33–35] transitions, complex plasmas constitute particulate systems with tunable behaviour [36], while the emergence of *de novo* proteins enables design of enzymatic behaviour [37].

Here we explore the possibility of tuning the collective behaviour in a biological system by changing the characteristics of the constituents in the bottom–up manner of the colloids and *de novo* proteins. For this purpose, we use the zebrafish system, which has a well-established collective behaviour that can be monitored in the laboratory by tracking the individual fish [39, 40, 77] and which is amenable to genetic modification [41–44]. Genetic modification has been shown to change behavioural patterns in groups of fish [45, 46]. By studying groups of mutant zebrafish, we can investigate how changes in individuals lead to collective behaviour and explore the underlying biological mechanisms that drive these emergent phenomena. The collective behaviour of zebrafish can further be compared to well established active matter models that encapsulate single fish properties [47–49], fish–fish interactions, [15, 20] as well as the interaction between fish and the environment [50, 51].

In particular, we investigated the behaviour of zebrafish carrying a *homozygous* deleterious mutation in the *col11a2* gene (*col11a2* -/-), that is to say, both parents exhibited the same mutation. These mutant fish exhibited abnormal skeletal development [52]. We also observed several degenerative defects in their vertebral column, including vertebral fusions and ruptures of the intervertebral discs. Therefore, we expect the mutation in *col11a2* to change the zebrafish swimming dynamics compared to the wildtype. Interestingly, we found that the *col11a2* mutant zebrafish exhibited limited vertebral column bending during swimming, and they took a significantly longer time to change their swim direction than wildtype. Surprisingly, a group of 25 mutant zebrafish exhibited a more polarised swimming pattern. These behavioral differences are well explained by the Vicsek model with a simple modification on the orientation updating rule [15, 40], suggesting that the *col11a2* mutant zebrafish exhibits reduced behavioral noise. Such reduction can be attributed to the limited bending of their bodies, intrinsically related to their vertebral defects. In this way, when interpreted through a simple active matter model, the state space accessed by the fish is controlled at the level of each individual in a manner analogous to simple tunable systems such as colloids.

## RESULTS

We bred genetically modified zebrafish whose *col11a2* gene is *knocked out*, such that the functional product of this gene is lacking, and compared the biological features of mutants to wildtype siblings. We found that the vertebral column of mutant fish displayed abnormal vertebrae. Additionally, they showed irregular shape of otoliths (mineralised rods located in the fish ears, which help them sense position), see supplementary Material (SM). Because these abnormalities could modify the dynamics of *col11a2 -/-* fish, we expect that the collective behaviour of the system may be modified at the level of single fish. To probe our hypothesis, we investigate the fish in two environments. Firstly, in a quasi–2d situation in which the fish are confined within a height of 5 cm, with a particular focus on their postures. We found that the *col11a2 -/-* zebrafish showed significantly reduced bending compared to the wildtype. Then, in a 3d swimming environment, we recovered the 3d trajectories of fish swimming alone (*N* = 1) or swimming as a group (*N* = 25) with a custom three view tracking apparatus [40, 53, 54]. Strikingly, a single mutant fish took a significantly longer time to change its orientation, and a group of *col11a2 -/-* zebrafish exhibited more ordered movement characterised by the polarization order parameter. By comparing our experimental data with numerical simulation of a modified Vicsek model, we found that the behavioral difference between mutant fish and wildtype fish is well captured by the model where on average *col11a2 -/-* zebrafish have lower orientational noise. Details of the fish tracking set-up are provided in the Methods and SM.

### The *col11a2 -/-* zebrafish exhibit defects in their vertebral column

To characterise the biological difference between the *col11a2 -/-* zebrafish and their corresponding wildtype controls (wt), we performed skeletal phenotyping to identify changes associated with bone and cartilage in the vertebral column. First, we performed microcomputed (X-ray) tomography (*µ*CT) of adult (6-month-old) zebrafish. which provides a 3d reconstruction of the fish skeleton. Strikingly, *col11a2 -/-* zebrafish (*n* = 16) commonly showed multiple vertebrae fusions associated with the caudal region of the axial skeleton. Such fusions compromised from 2 to up to 5 adjacent vertebrae, as shown in Fig. 1 (a). Using Alizarin Red staining, which labels calcified tissues and allows a detailed visualisation of the small zebrafish bones, we further detected bone abnormalities associated with the vertebral bodies and vertebral arches, exhibited in Fig. 1 (b).

**FIG. 1.**
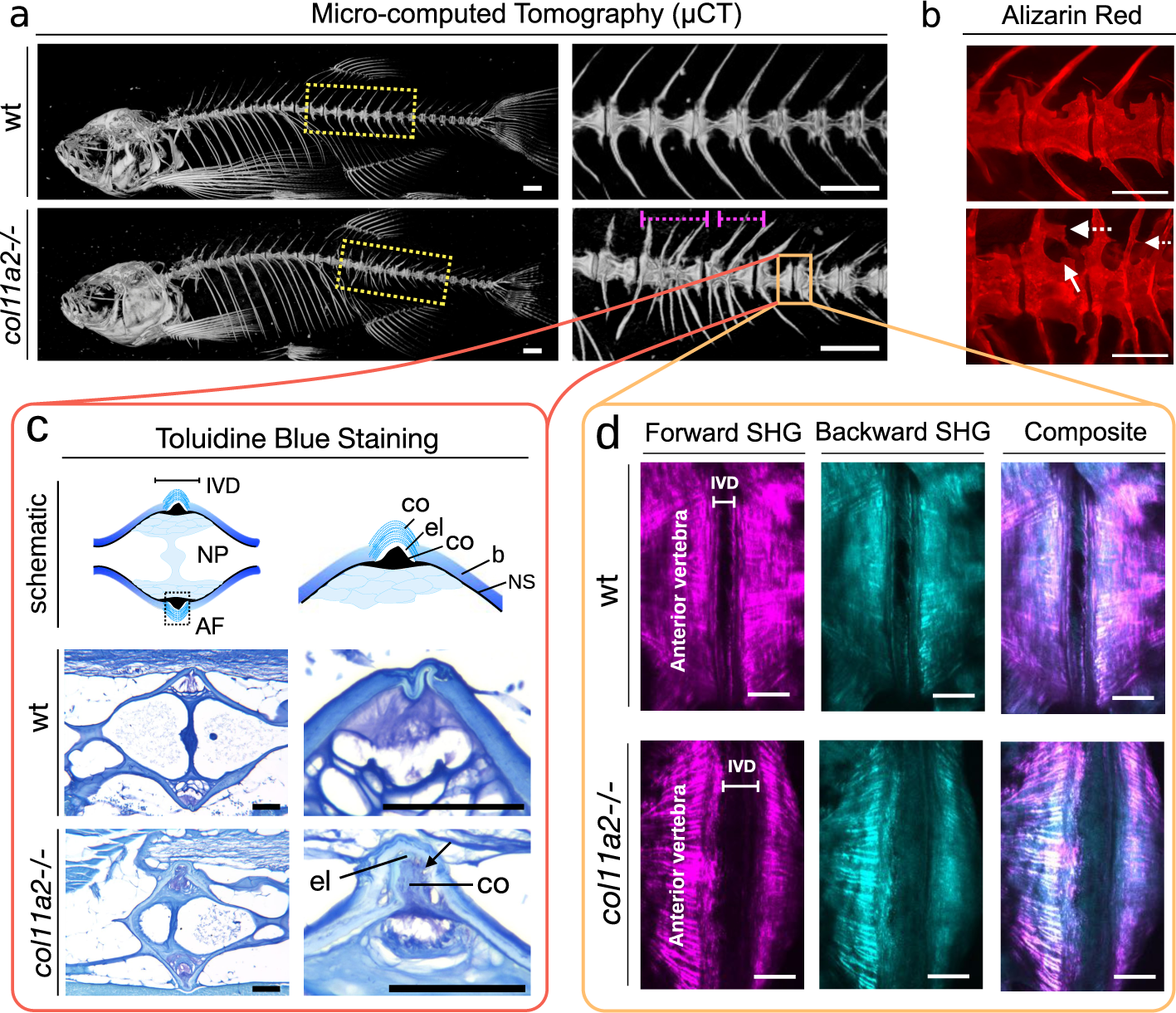
Effects of *col11a2-/-* knockout in zebrafish. The *col11a2 -/-* zebrafish have a fused vertebral column, ruptured annulus fibrosus and morphological abnormalities. (a) Micro-computed Tomography (*µ*CT) of 6-month wt and mutant fish. The region highlighted with dashed yellow box is show with higher magnification on the right panel. Note vertebrae fusions (magenta dash lines). (b) Fluorescent microscopic images of the calcified tissues of zebrafish labelled by Alizarin Red staining. Note abnormal vertebral trabeculations (arrow) and bone outgrowths from the arches (dashed arrow). Scale bars = 500 *µ*m. (c) Toluidine blue stained images showing the cellular components of the soft tissue intervertebral disc (IVD) region. The top schematic (left) shows the encounter of two vertebrae at the intervertebral disc (IVD) region, with the annulus fibrosus (AF) and nucleus pulposus (NP) annotated. The top right schematic shows a higher magnification view of the annulus fibrosus (AF) region, which is formed by an external layer of collagen fibres (co), an elastin layer (el) and another collagen rich region (co). The vertebral bone (b) and the notochord sheath (ns) are annotated. The sections of wt and *col11a2 -/-* fish are shown in the middle and bottom rows respectively. Note that the elastin (el) and the collagen (co) layers are ruptured (arrow). (d) Forward and backwards Second Harmonic Generation (SHG) images of wt and *col11a2 -/-* zebrafish intervertebral disc area. Note thicker collagen fibres at the endplates (dashed line) of mutant fish. Disorganised collagen fibres in the IVD (AF) is detected in mutants (arrowhead) Scale bars = 50*µ*m.

Intervertebral discs (IVDs) that separate one vertebrae from each other are the most important vertebral element to allow spine flexibility and serve as shock-absorbing cushions under mechanical load and vertebral column movement [55]. In mammals and fish, the IVD is formed by the nucleus pulposus (NP) internally (formed by fibrous tissue and cartilage-like cells) and externally at the bone junction by the annulus fibrosus (AF) composed of collagen and elastin layers [44] (Figure 1c). To assess the structure of the IVD at the cellular level, we imaged sections of wt and mutant fish, which were stained with by Toluidine Blue (which labels soft tissue), as shown in Fig. 1c. Here, tissue slices of 8*µ*m in thickness throughout the vertebral column were imaged. The regions forming the IVD are shown in the schematics. At lower magnification, the AF and NP can be observed (left panel). The NP is formed by cartilage-like cells and fluid. While the AF is formed by layers of elastin and collagen fibers. At higher magnification with a focus on the AF (right panel), the layers of collagen and elastin that form the AF can be observed in detail. These regions are well defined in the wild type fish but are very disorganized in the mutants. Strikingly, the higher resolution images (Fig. 1c, right) show a region of elastin that is ruptured in the mutant sample (indicated with an arrow). In other words, the most dramatic changes found in *col11a2 -/-* fish were associated with the annulus fibrosus (Fig. 1c). The collagen layers that compose the AF are abnormal, and the elastin layer, which allows flexibility to the IVD and the vertebral column, is ruptured. The NP is more fibrous than normal; however, these differences are not dramatic. Our results demonstrate that the lack of *col11a2* leads to a destabilization of the the IVDs and the elasticity of the vertebral column.

To better visualize collagen fibers associated with the AF, we imaged the fish vertebral column using second harmonic generation (SHG) microscopy. SHG imaging is a non-linear infra-red microscopy technique which is sensitive to molecular alignment, and therefore provides a method to examine the 3d architecture of the fibrillary collagen matrix [44, 56, 57]. Forward (transmission) mode and backward (backscattered) SHG are shown in Fig. 1d, which reveal significant changes in collagen organization at the verebral endplates (edges of the bones at the vertebra) of mutants. Specifically, we detected uneven endplates associated with increased thicker fibers of collagen at the AF interface. While in the AF region, disorganized thin collagen fibers were detected (Fig. 1d). Such dramatic changes in the organization of collagen fibers likely add to the flexibility impairment of the vertebral column during bending.

Our morphological assessment of wt and *col11a2 -/-* zebrafish from 3D *µ*CT images revealed another biological difference, an irregular morphology for the otoliths (small calcified rod-like structures inside the inner ear of the fish), presented in the SM. Otoliths are important in the regulation of balance during swim. Together, these observations characterise the structural difference of skeletons between mutant fish and wildtype, strongly supporting the biological foundation for the observed behavioral differences.

### Quasi 2d swimming: mutants have limited vertebral column bending

To understand the impact that the vertebral column had on swimming behavior, we analysed their swimming posture by placing individual zebrafish in the quasi 2d environment and used a camera above to capture videos while they swim, as illustrated in Fig. 2 (a) and (b). In these videos, the fish appear as an elongated ellipse that bends every now and then, as illustrated in the sequence of images in Fig. 2 (c). From our videos, we represent the shape of the fish with an image of *w* by *w* pixels, and flatten it into a *w*^2^ dimensional vector for each of the *m* images. In this way, we obtain a dataset of shape **X** ∈ *R^m×w^*^2^, and performed principle component analysis (PCA) to extract the characteristic modes of the fish shape. Specifically, we calculated the covariance matrix **C** = **X***^T^***X**, and then calculated all the eigenvectors of **C**. Typically, the eigenvector corresponding to the second largest eigenvalue of the covariance matrix describes the bending of the fish, and we term this eigenvector the *bending mode* **b**. We can, therefore, gauge the level of bending by projecting the fish shape (**x** ∈ *R^w^*^2^) onto the bending mode **b** by simply taking the dot product, **x** · **b**. This quantity is termed the *bending index*. A more detailed description of the PCA analysis and the visualisation of different modes are available in the SM.

**FIG. 2.**
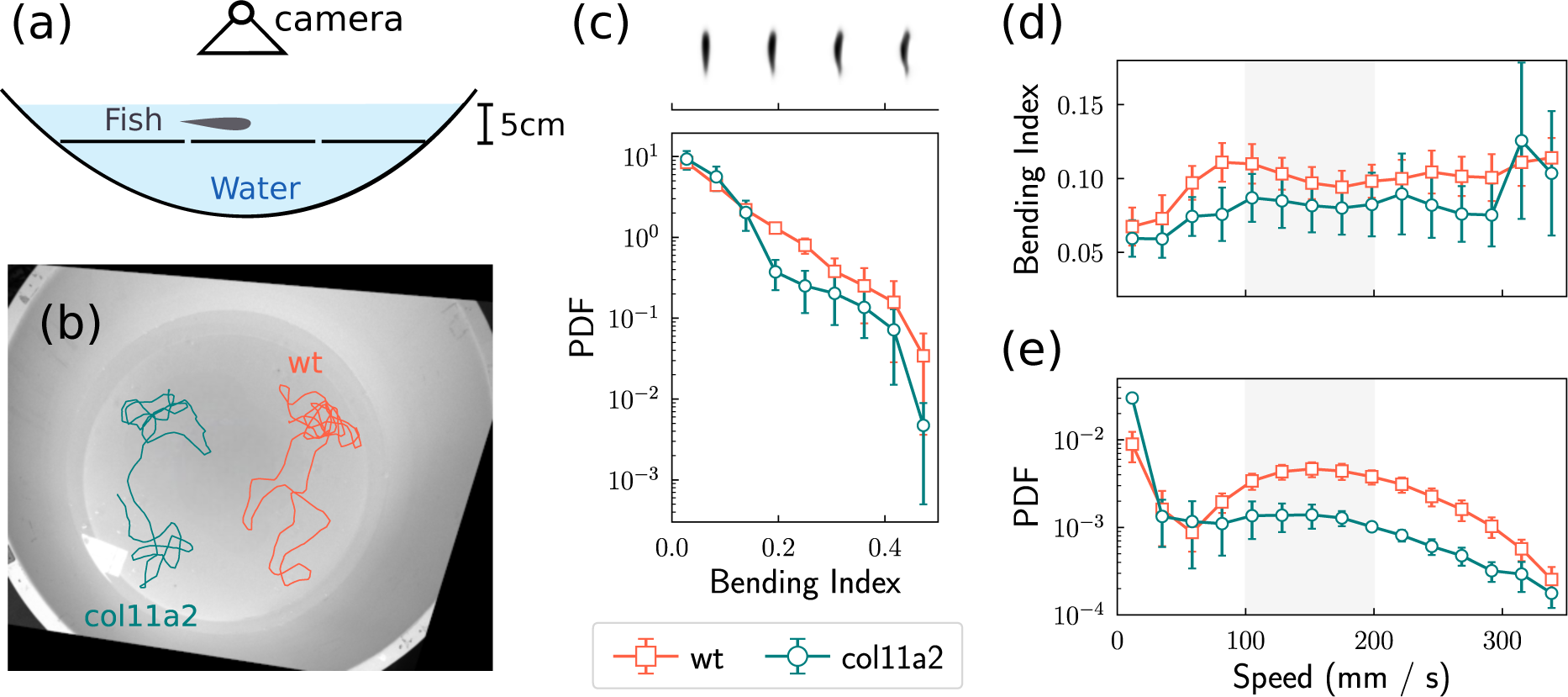
Differential swimming shape analysis of *col11a2 -/-* zebrafish. (a) illustration of our experimental setup for recording the 2d movement of fish. (b) Typical trajectories of mutant fish and wt fish replotted on top of the captured image. (c) Distribution of the bending index calculated from the image of the fish. The top panel, sharing the same x-axis with respect to the distribution plot, shows the characteristic shapes at different bending index values. (d) Average bending indices, calculated from the images of the fish shapes, for the wt fish and the mutant fish at different speed values. (e) Distribution of the speed values. Error bars represent the standard error calculated from 5 different fish.

The resulting posture analysis is shown in Fig. 2 (c) – (e) where we focus on the bending index of the mutant fish and the wildtype fish. The mutant fish exhibit less bending compared to the wildtype fish, as shown in Fig. 2 (c), indicated by the broader distribution of the bending index for the wildtype fish, as a result of “burst-and-coast” swimming style typical of zebrafish [58]. Recalling the discussion of Fig. 1 above, it is also likely that the mutant joints are “stiffer” than those of the wt, due to the defective elastin layer of intervertebral discs, so they favor swimming in a way that involves less bending.

The relationship between the bending index and the swimming of the fish is also informative, as plotted in Figs. 2 (d) and (e). The bending index of the wt fish reaches a maximum when they swim at speed around 100 mms*^−^*^1^, exhibiting decreased bending as their speed further increased from 100 mms*^−^*^1^ to 200 mms*^−^*^1^. This result explains the speed distribution of the wt fish, typically the bimodal shape that revealed a preferred swimming speed at around 150 mms*^−^*^1^, shown in Fig. 2 (e). Such a speed is preferred, possibly because in this speed range, they bend less and therefore use less energy. For the mutant fish, the correlation between the bending index and the swimming speed is rather different. The mutant fish exhibit smaller bending indices in speed regions from 0 to 300 mms*^−^*^1^, and their bending index varies less with respect to the speed, compared to the wt fish. The observed bending index of mutant fish also explained their speed distribution, particularly the lack of a bimodal shape. Specifically, the mutant fish do not have the swimming posture, that allows them to enjoy a higher speed while performing less bending, therefore, they do not have a preferred swimming speed. Importantly, we did not observe any behaviour changes that suggested an altered balance of the fish during the swim. Collectively, regardless of the various possible effects on the collective behavior associated with the mutants, we concluded that the vertebral column abnormality is the major contributor to behavior changes leading to reduced flexibility in mutant fish, which resulted in limited bending while swimming.

**3d swimming: Mutants take longer to change their swimming direction.**

This characteristic difference in speed distribution for the mutant fish and the corresponding wildtype, can also be observed when the fish swim in a 3d environment. We used a three–view apparatus to record zebrafish movement in 3d, and calculated individual 3d trajectories, as previously described [40]. An illustrative sketch of the apparatus is shown in Fig. 3 (a), with representative trajectories of mutant fish and wt shown in Fig. 3 (b). Specifically, we calculated 10 single–fish trajectories for both mutant fish and wt fish. We repeated the experiments with two different batches of fish, so that the batches are bred independently. Details of the experiments are available in the SM.

**FIG. 3.**
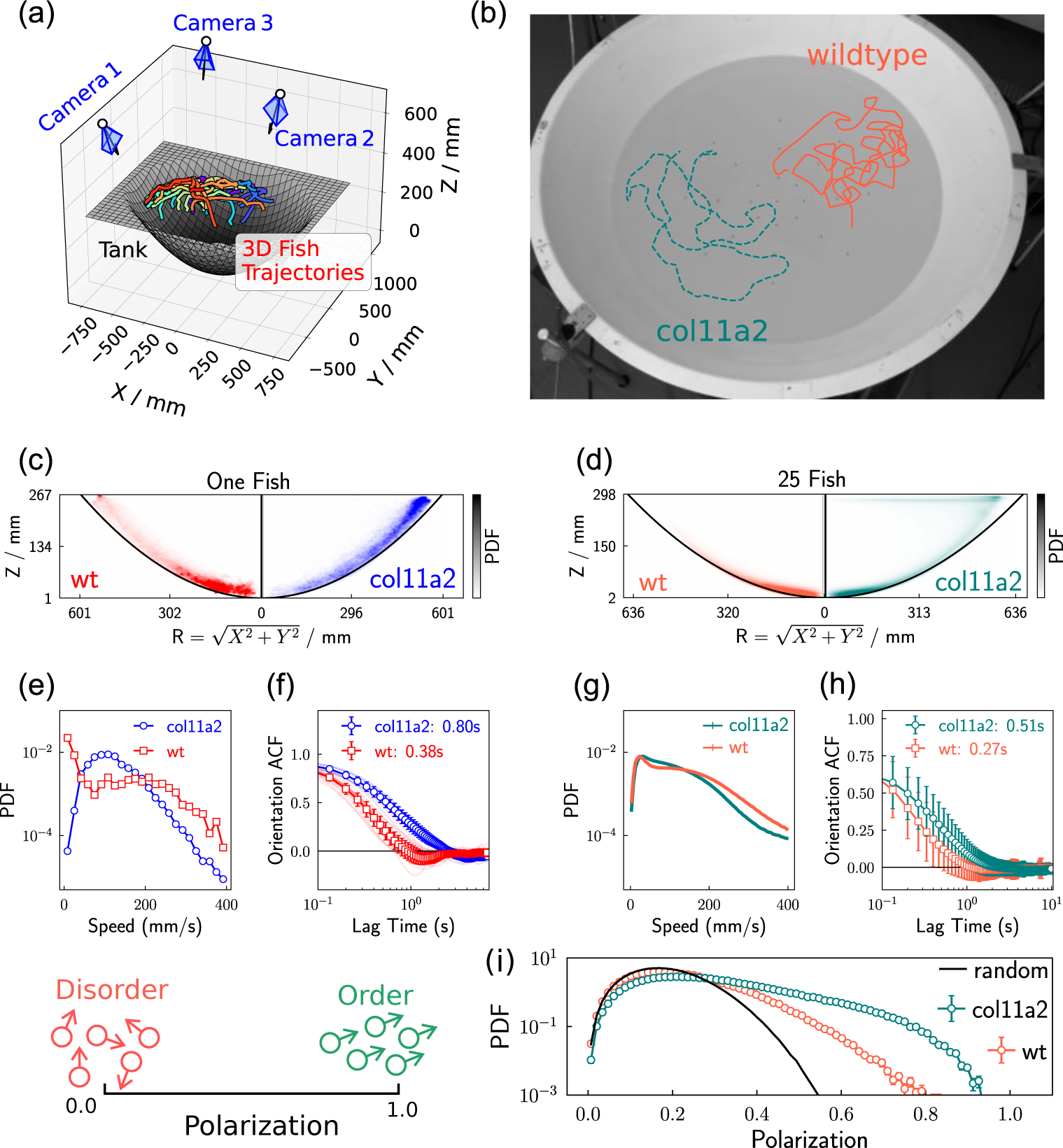
The behaviour of wt fish and mutant fish swimming in a three dimensional space. (a) A schematic of the experimental apparatus. The fish were placed in a bowl-shaped tank, and their movements were recorded by three synchronized cameras. The 3d trajectories of the fish were calculated from the recorded videos. The coloured line plots represent the movement of 25 mutant fish in 5 seconds. (b) The swimming trajectories for a single wt and a single mutant fish re–projected onto the recorded image, highlighting the difference in the persistence of the motion. (c, d) The joint probability distribution of the latitude radius (*R*) and the height (*Z*) of the wt fish and the mutant fish. (e, g) The probability distribution of the average swimming speed. (f, h) The averaged auto-correlation function of the fish orientation. (i) The probability distribution of the polarization order parameter (Eq. 1), whose definition is illustrated on the left. The solid black line represents the polarization distribution of 25 randomly orientated vectors.

The spatial distribution of individual wildtype and mutant fish is shown in Fig. 3 (c). Individual fish tend to move near the wall of the the tank, as seen in Fig. 3 (c) while few fish are distributed far from the tank wall. While the wt fish preferentially swim in the bottom region of the tank, the mutant fish are more likely to appear near the surface of the water. The wt exhibited their characteristic bimodal speed distribution in 3d, as shown in Fig. 3 (e), while the mutant fish exhibited a single peak around 100 mms*^−^*^1^ in their speed distribution. Such a difference is consistent with the 2d experimental results (Fig. 2 (c)), suggesting the mutant fish also bend differently while swimming in 3d. (The direct measurement of body bending in 3d is beyond our current capacity, but it is in principle possible [59].)

We now characterise the dynamics of the fish in a quantitative manner, using the orientational autcorrelation function (ACF). Here we detect significant differences in the dynamics of orientation between mutant fish and the wt fish. The mutant fish took longer to change their direction, indicated by the auto–correlation function (ACF) of the moving direction of the fish, as shown in Fig. 3 (e). The ACF depicts the temporal correlation of a random process at different time intervals, serving as a useful to calculate relaxation time scales from experiments and simulations [60].

Our analysis of zebrafish reorientation can be affected by several factors. For instance, the directional change of the non-moving individuals will be dominated by the tracking error. In addition, an individual in the bottom of the tank will be forced to change direction more frequently, because the otherwise ballistic motion will be interrupted by the tank. To exclude these extrinsic effects, we excluded the non-moving time points in the trajectories (speed *<* 50 mms*^−^*^1^), and focused only on a specific height region (50 mm*< z <* 150 mm) during the calculation of the ACF in Fig. 3 (f). To obtain the characteristic reorientation timescales (*τ*) of wt (*τ* = 0.38 ± 0.02 s) and mutant fish (*τ* = 0.80 ± 0.05 s), ACFs were fitted with an exponential function, *C*(*t*) = *A* exp(−*t/τ*) where *A*(≤ 1) is a constant. Our results thus show that mutant fish take longer to change their direction of movement than wt. The difference in the reorientation timescale was robust across successive measurements, as we repeated the experiment using zebrafish bred independently and obtained same results (see the SM for details on the experiments).

### A group of mutant fish exhibit more ordered motion

How does the change in the individual swimming pattern affect the behaviour of a group? To answer this question, we recorded the collective motion of 25 zebrafish in the same 3d tracking apparatus as that shown in Fig. 3 (a). We studied the fish at a variety of ages from 40 dpf (days post fertilisation) to 120 dpf, to capture the effects of fish development [61] and repeated exposure to the experimental environment [62]. This way, we can probe the swimming behaviour at different developmental times of zebrafish.

Even if we completely discard these developmental effects, by concatenating all the results together, we still find a significant difference between the mutant fish group and the wt group. The analysis of such a merged dataset is shown in Fig. 3 (d), (g) and (h). These differences are similar to those observed in the one–fish experiments. Again wt fish tended to be distributed at the bottom of the tank, exhibiting a bimodal speed distribution. When calculating the average orientational relaxation time (*τ*) of individual fish in the group, the mutant fish group also presents a higher value (*τ* = 0.33 ± 0.04*s*), compared to the wt group (*τ* = 0.20 ± 0.01*s*). The difference between a group of wt fish and the mutant fish is more significant if we compare the two groups in a specific age, as shown in the supplementary material.

Strikingly, groups of mutant fish exhibit more ordered motion, as shown in Fig. 3 (i). The degree of order of the motion is characterised by the *polarization* (Φ), a dynamical order parameter with the following definition,

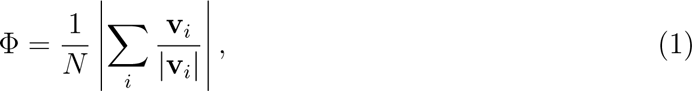

where *N* is the number of fish in the group, and **v***_i_* represents the velocity of the *i*th fish in the group. The value of Φ approaches unity if all the fish swim in the same direction and zero in the case that the swim directions are random. The distribution of Φ for mutant fish is significantly shifted towards larger values, suggesting that the collective behaviour of mutant fish is more ordered. This difference is visually obvious as shown in the supplementary video (sMovie01.mp4).

### The collective behaviour of mutants can be modelled by a modified Vicsek model

The increase in ordering in the case of the *col11a2 -/-* zebrafish, compared to the wildtype may be explored in the context of the Vicsek model [15], which exhibits an order to disorder transition with the increase of orientational randomness or the decrease of number density. The Vicsek model is a minimal model where the simulated agents only interact with each other via local orientational alignment, and this model can be modified to match the fish behaviour, simply by adding an inertial term such that the velocity **v***_i_* of each agent (fish) is updated as, [27]

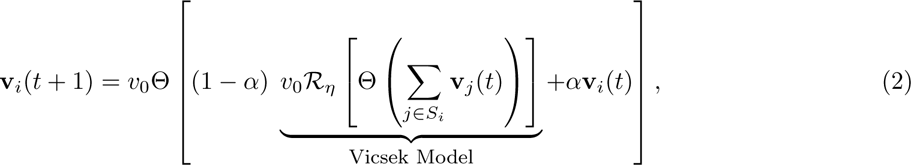

where *v*_0_ represents the constant self–propelling speed for all agents. Θ normalises the velocity vector, such that the constant velocity condition is maintained. Noise in the direction of motion is implemented with *R_η_* such that the velocity vector falls on a spherical cap of amplitude 4*πη* with *η* controlling the strength of the noise [27]. The parameter *α* takes the value between 0 and 1, and effectively adjusts the inertia of the agents, so that those with larger *α* values change their orientations more slowly. Essentially, the velocities for all agents in time point *t* + 1, **v**(*t* + 1), are mixtures of the Vicsek updating results, as well as their respective velocities in time point *t*, **v**(*t*). Such “memory” in terms of velocities gives the agents effective inertia in their orientations, which prevents the agents from performing unrealistically fast reorientations. The value of *α* in Eq. 2 changes the relative proportion of the two, and we set its value to be 0.63, consistent with previous work [40]. In our discussion on the effect of the *α* value of the model is available in the, we conclude that the *α* = 0.63 fitted to our data provides convincing evidence for some effective inertia in the zebrafish system.

The model generates 3d trajectories that can be compared to the experimental data. Such comparison can be made easy if we measure dimensionless quantities from both simulations and experiments, as we can therefore ignore biological details such as the body length of the fish. Here we focus on the *reduced persistence length*, defined as the ratio between the persistence length *l_p_*, and the average nearest neighbor distance *l*_nn_. The persistence length *l_p_* is the product of the orientation relaxation time *τ* and the speed of the fish/agents, averaged between all individuals. The value of *l*_nn_ is calculated by averaging the distances of all fish/agents to their closest neighbors. Both *l_p_* and *l*_nn_ have units of length so that their ratio is dimensionless. With these measures in hand, we now proceed to investigate whether the *col11a2 -/-* zebrafish may similarly be described with the same model.

In Fig. 4(a), we plot the values of persistence length (*l_p_*) and nearest neighbour distance (*l*_nn_) for both the mutant (squares) and wildtype (circles) fish. We see that the mutant fish have both larger *l*_nn_ and *l_p_*values. The larger *l*_nn_ values of the mutant fish reflect their distribution in the upper location of the tank, of naturally larger volume given the geometry of the tank. The larger *l_p_* of the mutant fish is presumably related to their slow re-orientation, as the persistence length *l_p_* is defined as the product of the speed and the orientational relaxation time. With the aging of the zebrafish, the average *l*_nn_ decreased for both the wt group and the mutant group, indicating that the fish exhibited more cohesive behaviour as they grew from 43 dpf to over 100 dpf. This increasing local density amongst zebrafish was also observed by Buske and Gerlai, and could be related to ecological factors such as avoiding predators, mating, or food discovery [63]. For juvenile fish (ages from 43 to 103 dpf), whether wt or mutant, a correlation between *l_p_/l*_nn_ and Φ was also observed, as shown in Fig. 4 (b). The mutant fish have a larger reduced persistence length compared with wt fish, which is correlated with higher polarisation.

**FIG. 4.**
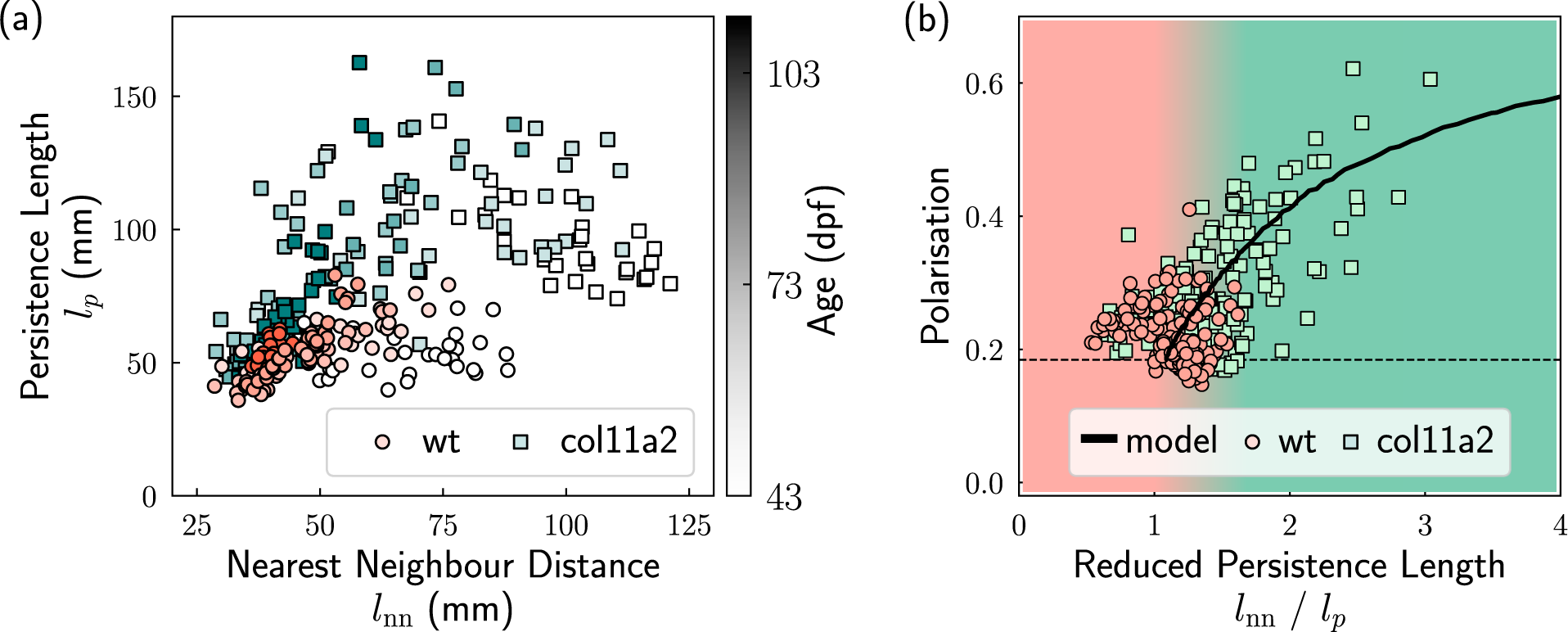
**Important lengthscales of 25 wt fish (**0**) and mutant fish (**D). (a) The nearest neighbour distance and the persistence length for the fish. Each scatter represents the average value in 2 minutes, whose grayscale values represent the age of the fish. (b) The relationship between the group polarisation and the reduced persistence length values. The data points represent the experiments and the line represents the simulation result where the fish obey the updating rule in Eq. 2. The shading represents the regimes of the reduced persistence length which are dominated by the wildtype (red) and mutant (green). The dashed line represents the expected polarization of 25 randomly orientated vectors.

The simulation results of the model provide a good fit for the experimental results, as shown in in Fig. 4 (b). To obtain different persistence length values, we changed the orientational noise of each individual agent. The match of the experimental data to the model suggests that the fish were effectively aligned with each other. Such alignment interaction enables the fish group to access an ordered state with high polarisation. The lack of *col11a2* caused defects in their spines, changed their swimming postures, and resulted in slower re–orientation of individuals, hence increasing the persistent length *l_p_* of the group. The increase of *l_p_* for a group of mutant fish yields more ordered movement for the presence of the aligning interaction. Nevertheless, at the level of our analysis, these changes in behaviour can be described with the same model we used previously.

In short, the same modified Vicsek model (with the same value for the inertia parameter *α*) fits both wildtype and *col11a2 -/-* zebrafish. In the context of this model, the mutants sample, on average, a different region of state space, although the distributions between the two types of fish do overlap. To the best of our knowledge, this is the first report of the explicit linkage between the macroscopic phenomena (group polarisation) of the animal collective behaviour, and the microscopic detail of the individual animal (spinal defects and changed swimming posture) which we control with genetic modification.

## DISCUSSION

Knocking out the *col11a2* gene changes the zebrafish system at the level of the individual fish, in several ways. The gene regulates the expression of the collagen, and its absence leads to defects in their vertebrae and intervertebral discs. Consequently, the mutant fish exhibit less bending during swimming compared to the wt fish, especially when the speed is slow. The compromised spinal development and bending ability are likely to prevent the mutant fish from making frequent directional changes when swimming, leading to a longer orientational relaxation time. Effectively, the *col11a2 -/-* zebrafish exhibited less “orientational noise”, if they were modelled as self–propelling agents with aligning interactions. On the group level, the reduced noise value is captured by the increased persistence length. A group of mutant fish is then expected to swim in a more ordered manner, where all the fish are likely to swim towards the same direction. This expectation is consistent with our observations for groups (*N* = 25) of wt fish and *col11a2 -/-* fish. In other words, we can use the genetic modification to tune the fish behaviour in a way which can be interpreted with a simple physical model.

The behaviour of animals in a group with moderate size, is affected by both the microscopic details of individuals and the interaction between individuals. In our study, the collective motion of 25 fish is affected by the microscopic, single–fish property, as shown in Fig. 3. Nevertheless, the fish–fish interaction also triggered a significant difference in the group polarization (Fig. 3 (i) and Fig. 4). For biological purposes, the microscopic single–fish features are useful measurements of the phenotypic response to the genetic modification. In this study, we followed this “biological path” extensively by looking into the single–fish behaviour as well as the bone structures of the fish. On the other hand, we expect the knockout of the *col11a2* gene to trigger a larger response in a larger group, since the discontinuous nature of the dynamical order–disorder transition [17, 28] will be blurred out in a small group, as a consequence of the finite-size effect [64]. However, performing such an experiment with a large group of mutant fish is challenging, as mutants display a relatively high mortality rate. In addition, tracking larger groups requires better camera systems (see [65] for example), more optimised algorithms (see [66, 67] for examples), and a very large space to avoid the interruption from the boundary. These requirements are beyond our current experimental capability. Nevertheless, we would expect a large group of wt fish to stay in the order-disorder transition region to enjoy the maximum susceptibility [40, 68, 69], while a large group of mutant fish are expected to reside on the ordered region.

## CONCLUSION

Our work describes a quantitative study on the behavioural difference between wildtype zebrafish and the *col11a2 -/-* zebrafish whose *col11a2* gene is knocked out (suppressed). The genetic modification caused defects in the vertebral column of mutants, leading to less body– bending in swimming, hence a slower reorientation process. This behavioural difference of a single fish affected the group behaviour of twenty five, where the mutant fish group exhibited more ordered collective motion that correlates with their more persistent swimming style. In other words, we have tuned the behaviour of a biological system in a manner reminiscent of synthetic colloids or *de novo* proteins. This is interpreted as the mutant fish system adopting a different state space with respect to the wildtype in the context of a minimalistic physical model. In particular the difference between mutant and wt fish can be summarised by different orientational noise level in a modified Vicsek model. Taken together we have shown that by modelling the swimming behaviour of zebrafish and treating them as active matter, we can reveal physiologically relevant behaviours that are underpinned by changes to the skeleton, demonstrating the utility of this approach as a tool to follow genetic mutants and relate behaviours to genetic traits and to phsyiological changes within the body.

## METHODS

### Zebrafish Husbandry

The wildtype zebrafish and the mutant fish were bred and kept in the fish facility in the university of Bristol, following standard procedures [70]. Our behaviour experiments were approved by the local ethics committee (University of Bristol Animal Welfare and Ethical Review Body, AWERB) with a UIN (university ethical approval identifier). *col11a2* sa18324 mutant zebrafish have a C *>* A base pair change at position 228, which leads to a premature stop codon the triple helical domain of the *α*2 chain of collagen 11.

### Micro-computed tomography (*µCT*) and bone density analysis

Adult zebrafish (6 and 12-month-old) were fixed in 4% PFA for one week and sequentially dehydrated to 70 % ethanol. After fixation, full body scans were taken with a voxel size of 20 *µ*m, using a Nikon XT H 225ST *µ*CT scanner (x-ray source of 130 kV and 53 *µ*A without additional filters). The generated scans were then reconstructed using Nikon CT Pro 3d software. During reconstruction, greyscale values were calibrated against a scan of a phantom with known density (0.75 gcm*^−^*^3^) using the exact same settings. Calibrations with the phantom were used to calculate bone density. Morphological analyses were performed using Avizo software (FEI) and volume rendering function. Tissue Mineral Density (TMD) was calculated based on the segmentation of the centra from caudal vertebrae (position 1-3).

### Alizarin Red Staining

Alizarin Red staining was performed on 1-year-old zebrafish, as previously described [71]. Pictures were taken using a Leica Stereomicroscope (DM4000) with a 580 excitation filter.

### Sections for Toluidine Blue imaging

Adult fish (1-year-old) were fixed in 4% paraformaldehyde (PFA) for 14 days, then decalcified in 1M Ethylenediaminetetraacetic acid (EDTA) solution for 20 days at room temperature. Samples were dehydrated to ethanol, embedded in paraffin, and sagittal sections at 8 µm thickness were taken. Selected slides were de-waxed and stained with Toluidine Blue [72]. Images were acquired on a Leica DMI600 inverted microscope, using 20X and 40X oil objectives, LAS software, and a DFC420C colour camera.

### Second Harmonic Generation (SHG)

Dissected spines of wt and *col11a2 -/-* zebrafish (1-year-old) were mounted in 1% low-melting point agarose (n = 3 for each group) and SHG imaged acquired using 25x/0.95 water dipping lens, 880nm laser excitation, and simultaneous forward and backward detection (440/20) with a Leica SP8 AOBS confocal laser scanning microscope attached to a Leica DM6000 upright epifluorescence microscope with multiphoton lasers allowing fluorescent and SHG acquisition of the same sample and z-stack. Microscope parameters for SHG acquisition were set as previously described [56].

### 2d Tracking Apparatus

We used one camera (Basler ac2040um) to capture the motion of a single fish at a rate of 15 frames per second. The camera was mounted on top of the fish tank, which offered a shallow (5 cm) water environment for the fish. Since it is challenging to mount camera perfectly perpendicular with respect to the water surface, we further rectified the obtained image following the method introduced in [53] by capturing an image with a chessboard floating on top of the water surface. We segmented the pixels corresponding to the single fish and took the centre as the 2d fish location. The locations in different time points were connected to form the trajectories.

### 3d Tracking Apparatus

We used three synchronised cameras (Balser ac2040 um) to capture the motion of the fish at a rate of 15 frames per second. The intrinsic parameter of the cameras was calibrated with a standard chessboard pattern. The extrinsic parameters, containing the location and orientation of the cameras, were calculated from a chessboard pattern floating on the water surface. The results were then optimised by enforcing a geometric constraint from the tank, whose shape is known.

The observation setup is composed of a white bowl-shaped container (referred to as the observation tank), which was immersed in a circular paddling pool. The water temperature of the observation tank was maintained around 25 *^◦^*C by heaters in the paddling pool, and we placed two filters in the paddling pool to control the water quality.

The fish were transferred from their living tank to the observation tank before each experiment. The video recording started 5 minutes after the transfer, and the room was evacuated for 5 minutes. The water in the observation tank was still during the observation, and we circulated the water when the fish were not being observed.

### Data Processing

We explicitly considered the projective geometry to calculate the 3d locations of the fish from their corresponding images from different cameras [53]. During the calculation, we considered the refraction of light at the water-air interface were considered, following our previous method [40]. We linked the 3d locations into short trajectories following a four-frame method [73], and extended these trajectories following the idea of Xu [74]. We take the time difference of locations as velocities and the directions of the velocities as the fish orientations. The autocorrelation function of the orientations was calculated to determine the reorientation time. The code for our image process and correlation analysis is available as an open-source project on GitHub (https://github.com/yangyushi/FishPy).

For the swimming posture analysis, we segmented the fish from the background in the image captured by the camera. The segmentation was performed by setting all pixels whose intensity values are below the threshold value to background pixels. We aligned the orientation of the fish by rotating the principle axes of the foreground pixels, so that these axes align with the X and Y axis of the image. The segmented and aligned image contain one fish, with the size of 50 × 50 pixels. We then concatenated all the 37,985 aligned images, from both the wt fish and the mutant fish across different observations in a quasi-2d environment, into a single array with a shape of 37, 985 × 50 × 50. The array was then flattened into shape 37, 985 × 2500 for the PCA. Effectively, we analysed a dataset with 2500 features and 37, 985 samples.

### Simulation Details

We simulated a variation of the Vicsek model proposed in [40] whose equation of motion is given by Eq. 2. To fit the experimental data to this inertial Vicsek Model, we changed the value of the noise parameter *η* from 0.5 to 1.0, while fixing the values of other variable (*v*_0_ = 0.1*, ρ* = 0.5*, α* = 0.63). These fixed parameters also fits the behaviour of 50 adult zebrafish, except for the value of the density (we set *ρ* = 1 for 50 adult fish). The consistent fit suggests our proposed model, regardless its simplicity, does capture the essence of the collective behaviour of zebrafish.

We use our custom Python library to carry out the simulation of the inertial Vicsek model. To fit our experimental results, we set the inertial term (*α*) to be 0.63, the number density (*ρ*) to be 0.5, and the speed (*v*_0_) to be 0.1. The agents were confined by a cubic box with periodic boundary conditions. We carried out simulations at different noise intensity values ranging from 0.65 (less noisy) to 1.0 (completely random). At each noise value, we prepared the system with random locations and random orientations. We waited for 10^5^ simulation steps for the system to reach a steady state and sampled another 10^5^ steps to collect data for statistical analysis. The simulation library is available as an open-source project on GitHub (https://github.com/yangyushi/FishSim).

### Data and Code Availability

Representative data is available here: https://zenodo.org/records/10693744. Simulation code is available on GitHub (https://github.com/yangyushi/FishSim).

## ACKNOWLEDGEMENTS

JL Ross Anderson is gratefully thanked for discussions. YY gratefully acknowledges the China Scholarship Council for funding, and YY thanks Min Wang for the help with editing the supplementary video. EK was funded by Versus Arthritis (Career Development Award, 23115). CPR acknowledges the European Research Council (ERC consolidator grant NANOPRS, project 617266).

